# Strange Stable Replicators Generated From Mumps Virus cDNA Clones

**DOI:** 10.1101/328179

**Authors:** Connor Bamford, Elizabeth Wignall-Fleming, Richard Randall, Paul Duprex, Bert Rima

**Affiliations:** Centre for Virus Research, Glasgow, UK; School of Biology, St Andrews University, UK; National Emerging Infectious Disease Laboratory Boston University, USA; Centre for Experimental Medicine, Queen’s University Belfast, UK

**Author notes:** **these authors contributed equally to the paper** (Connor Bamford, Elizabeth Wignall-Fleming).

**Keywords:** mumps virus, mutation, cluster, next generation sequencing, genetic stability, quantum effects

## Abstract

In reverse genetic experiments we have isolated recombinant mumps viruses (rMuV) based on a recent clinical isolate that carry large numbers of mutations clustered in small parts of their genome and which are not caused by biased hyper-mutation. In two separate experiments we obtained such rMuV: one virus had 19 mutations in the V/P region of the genome; the other, which also contained an extra transcription unit encoding green fluorescent protein (EGFP), had 32 mutations in the N gene. These specific constellations of mutations have not been observed in naturally occurring MuV isolates. The vast majority of the mutations (48/51) are synonymous.

On passage in Vero cells and human B-LCL cells, a B lymphocyte-like cell Line, these mutations appear stable as no reversal occurs to the original consensus sequences, though mutations in other genes occur and change in frequency during passage. Defective Interfering RNAs accumulate in passage in Vero cells but not in B-LCL cells. Interestingly, in all passaged samples the level of variation in the EGFP gene is the same as in the viral genes, though it is unlikely that this gene is under any functionality constraint. The stability in repeated high multiplicity passage indicates that the constellation of mutations is placing the virus on a fitness peak from which it cannot escape. What mechanism gave rise to these mutant viruses and their stability remain open questions of interest to a wider field than mumps reverse genetics alone.

## Introduction

Mumps virus (MuV) is a human pathogenic RNA virus in the genus *Rubulavirus* in the family *Paramyxoviridae* (Rubin, Sauder and Carbone, no date). This family as non-segmented negative stranded RNA viruses shares basic replication strategies with the other viruses in the order *Mononegavirales*. MuV has a genome of 15,384 nucleotides (nt) in length, which contains 7 transcription units from the 3’ end of the negative stranded genome to the 5’ end respectively. These encode respectively the nucleocapsid protein (N); the innate immune modulatory protein V, the matrix protein (M), the fusion protein (F), a small hydrophobic protein (SH), a haemagglutinin-neuraminidase protein (HN) and the large protein (L) which carries the RNA-dependent RNA polymerase activity (RdRp). The major co-factor for the RdRp required during transcription is the phosphoprotein (P) which is generated by co-transcriptional editing during the copying of the V gene to generate P protein by the insertion of 2 (or 5) non-templated G residues into the nascent transcript. A so-called I protein is also generated by insertion of 1 or 4 G residues. V and I are non-structural proteins. In the *Rubulavirus* genus there is no overlapping open reading frame (ORF) for the C protein in the gene encoding the V and P proteins and unedited mRNAs transcribed from the MuV genome encode the V protein and not the P protein as is the case in many of the other *Paramyxoviridae* (Rubin, Sauder and Carbone, 2013). No overlapping ORFs have been identified in the N gene of MuV.

Reverse genetics of MuV and other members of the order *Mononegavirales* from plasmids that encode the entire genome of the virus has been described for over two decades now (Clarke *et al.*, 2000)and this system has been used to elucidate aspects of mumps virus pathogenesis and virulence (Rubin, Pletnikov and Carbone, 1998; Clarke *et al.*, 2000; Lemon *et al.*, 2007; Sauder *et al.*, 2011; Xu *et al.*, 2012). In this, it is no different from other viruses in the order *Mononegavirales.* Recently, we established a ‘rescue’ system based on the sequence of MuV in clinical tissue material so that we could study the properties and behaviour of viruses that had not been passaged in cultured cells and thereby potentially exposed to selective pressures exerted by the host cell *in vitro*. Viruses were rescued on Vero cells (MuV^G09^) from the clinical material and proved to be genotype G virus, similar to other viruses isolated in the US during outbreaks (Xu *et al.*, 2011). Details of the virus and its rescue and use in the study of mumps pathogenesis will be described elsewhere.

MuV is stable in the field and replacement rates have been estimated to be approximately 3 × 10^-4^ per annum (Pomeroy, Bjørnstad and Holmes, 2008). No laboratory estimate for mutation rates has been reported. Here, we report that we obtained a number of rescued viruses that carried clusters of mutations in the N and V/P genes, which involved stable synonymous nucleotide changes that have not been observed in natural isolates in any of the genotypes of MuV hitherto described.

## Results

### Generation of recombinant MuV with clusters of mutations

Our attempts to obtain rMuV virus failed in all cases to give us a virus with the same sequence to that in the plasmid after aspirating the material of a primary syncytium in a single well and passaging this 4 times on Vero cells at low MOI. The reverse genetics system used ‘helper’ expression plasmids representing the authentic N, P and L genes of MuV^G09^ and a plasmid containing the consensus sequence of MuV^G09^ in order to prevent the occurrence of potential recombination events between the helper plasmid and the full length genome plasmid. Out of the 5 plaque picks that were made in the primary rescue wells transfected with a plasmid representing the full length genome of MuV^G09^ none were identical to the input cDNA clone sequence and a combined total of 18 mutations were observed; one virus rMuV^G09^PP1 (abbreviated as PP1) had 11 mutations in the V/P gene (Table 1) between nucleotides 2551 and 2867 spacing mutations on the average 32 nt from each other. This area encodes the C terminal end of the V protein and part of the more conserved C-terminal domain of the P protein in MuV. Nine mutations were synonymous and two of the PP1 mutations led to amino acid changes at position 192 (P>L) and 212 (Q>P) in the P protein. Both mutations were synonymous in the overlapping V protein reading frame.

**Table 1:** Stability of the unique mutations in MuV-PP1 after 6 passages on Vero cells.

| Position<br>nr in the<br>genome | Change<br>from G09<br>to PP1 | Effect in PP1 | Reads in PP1<br>passage 6 | Variant<br>reads | Reads in<br>MuV<sup>G09</sup><br>passage 6 | Variant<br>reads |
| --- | --- | --- | --- | --- | --- | --- |
| 2551 | C=U | P192P=L;<br>V191P=P syn | 276 U | none | 1606 C | 1 A |
| 2611 | A=C | P212Q=P<br>V211P=P syn | 207 C | **1 A**; 2 U | 1171 A | 1 A; 1 C |
| 2654 | A=G | P226R=R syn | 206 G | **3 A** | 986 A | none |
| 2763 | C=U | P263L=L syn | 222 U | none | 1544 C | 1 U |
| 2768 | U=A | P264A=A syn | 232 A | none | 1312 U | 3 C; 1 G |
| 2780 | A=G | P268G=G syn | 206 G | none | 1417 A | none |
| 2789 | G=A | P271A=A syn | 265 A | **1 G** | 1635 G | 1 U |
| 2810 | G=U | P278P=P syn | 261 U | none | 2149 G | 2 U |
| 2816 | C=U | P280N=N syn | 266 U | 1 A; 2 G | 1645 C | 2 A |
| 2864 | U=C | P296H=H syn | 280 C | none | 1714 U | none |
| 2867 | A=G | P297V=V syn | 294 G | none | 1609 A | none |
| Total |  |  | 2715 | 10 | 16788 | frequency<br>0.8 x 10<sup>-3</sup> |

Syn= synonymous; in bold variants that would restore the original nucleotide in MuV<sup>G09</sup>

Rescue was also attempted from a plasmid into which the EGFP gene had been inserted between the V/P and M genes of mumps virus MuV^G09^. Similarly, out of the 7 plaque picked viruses none had a nucleotide sequence identical to the original plasmid. Six out of the 7 viruses carried *in toto* 9 mutations (8 non-synonymous replacements and one insertion in the poly-adenylation signal in the F gene). One plaque picked virus rMuV^G09^EGFP(3)PP2 virus (abbreviated here as PP2) had 32 mutations in two clusters in the N gene (Table 2). One cluster contained 13 mutations between nt 607 and 860 distancing mutations by on average 21 nucleotides. This area encodes a relatively conserved part of the N protein of MuV. The second cluster contained 19 mutations between nucleotides 1225 and 1558 with an average spacing of 18 nucleotides. This encodes part of the relatively variable C terminal tail of the N protein of MuV. In PP2 almost all the mutations (31/32) were synonymous with the exception of a single V>A mutation at position 460 of the N protein. None of the mutations observed in PP1 and PP2 were observable as minor peaks in sequencing chromatograms of the sequencing reactions carried out on the clinical material. The constellations of synonymous mutations observed in these viruses in the N and V/P genes were also not present as linked variations in sequence of the many mumps virus genotypes in the databanks. No clusters of mutations were observed in the other genes of MuV by Sanger sequencing of the primary isolates.

**Table 2:** Stability of the unique mutations in PP2 after 6 passages on B-LCL cells.

| Position<br>nr in the<br>genome | Change<br>from G09<br>to PP2 | antigenome<br>reads in<br>passage 6 | Variant reads | genome<br>reads in<br>passage 6 | Variant reads |
| --- | --- | --- | --- | --- | --- |
| 607 | G > A syn | 846 A | 1 C | 850 A | 2 C |
| 622 | A > G syn | 1031 G | none | 1001 G | none |
| 628 | A > G syn | 1067 G | 4 U | 1059 G | none |
| 679 | C > A syn | 680 A | none | 1506 A | 1 G; 1 U; **1 C** |
| 697 | U > G syn | 692 G | none | 2502 G | **3 U** |
| 736 | G > U syn | 887 U | 5 C; **2 G** | 2733 U | 6 A; 1 C; **7 G** |
| 760 | G > U syn | 1093 U | 4 C; **3 G** | 1594 U | **1 G** |
| 784 | A > G syn | 975 G | **3 A** | 1348 G | 6 U; **2 A** |
| 841 | U > C syn | 1338 C | **2 U**; 2 A; 2 G | 659 C | 2 A |
| 844 | U > C syn | 1400 C | 3 A; **3 U** | 650 C | 2 A; 1 G |
| 850 | U > C syn | 1447 C | **1 U**; 1 G | 632 C | 1 G; 1 U |
| 859 | A > G syn | 1444 G | 2 U | 523 G | none |
| 860 | C > U syn | 1434 U | **3 C**; 3 G | 489 U | none |
| Total<br>frequency | cluster 1 | 14334 | 43 | 15546 | 37<br>1.04 x 10<sup>-3</sup> |
| 1225 | C > U syn | 1325 U | **2 C** | 620 U | **1 C**; 1 G |
| 1240 | G > A syn | 1461 A | 6 C; 1 U | 514 A | 1 C |
| 1312 | A > G syn | 2819 G | 2 U; **1 A** | 369 G | 2 U |
| 1327 | C > U syn | 2613 U | **6 C**; 2 G; 1 A | 339 U | **1 C** |
| 1330 | U > C syn | 2645 C | 10 A; 8 U; 1 G | 343 C | **1 U** |
| 1354 | A > G syn | 2759 G | **12 A**; 2 C; 4 U | 488 G | none |
| 1366 | G > A syn | 2811 A | **5 G**; 2 C; 1 U | 764 A | **3 G**; 1 U |
| 1369 | A > G syn | 2836 G | 9 U; **3 A**; 2 C | 784 G | 1 C; 1 U |
| 1384 | G > C syn | 2596 C | 6 A; 3 U | 1195 C | 3 A; 1 U |
| 1387 | G > A syn | 2610 A | 8 C; 2 U; **1 G** | 1322 A | 1 U] |
| 1390 | C > U syn | 2584 U | 4 A; 2 G | 1429 U | 3 A; 3 G |
| 1413 | G > A syn | 3626 A | 3 C; 2 U; **1 G** | 1612 A | 3 U; 1 C |
| 1438 | G > A syn | 3749 A | 6 C; **5 G**; 3 U | 1553 A | 8 C; 3 U; **1 G** |
| 1451 | U > C syn | 3508 C | 5 A; **3 U**; 2 G | 1510 C | 3 A; **1 U** |
| 1468 | C > U syn | 4457 U | **5 C**; 3 G; 2 A | 982 U | **4 C**; 2 G; 1 A |
| 1471 | U > C syn | 4496 C | 3 A; **2 U** | 1167 C | 2A; **1 U** |
| 1489 | U > C syn | 4259 C | **7 U**; 5 A | 1352 C | **2 U** |
| 1524 | U > C<br>N460V=A | 4359 C | 5 A; **3 U** | 1265 C | 4 A; 1 G; **1 U** |
| 1558 | G > A syn | 2910 A | 3 U; 2 C; **1 G** | 729 A | **1 G**; 1 U; 1 C |
| Total<br>frequency | cluster 2 | 58423 | 176 | 18337 | 65<br>1.05 x 10<sup>-3</sup> |
| Total | both<br>clusters | 72757 | To wt: 81 other:<br>138<br>Other/wt<br>= 1.70 | 33883 | To wt: 31<br>To other: 71<br>other/wt =<br>2.29 |

In bold variants that would restore the original nucleotide in MuV<sup>G09</sup>

The original mutations first determined by Sanger sequencing in the V/P gene of PP1 and the N gene of PP21 were confirmed in Next Generation Sequencing (NGS) studies described below, with the exception of a deletion mutation in the L gene in the recombinant PP1 virus. In PP1, Sanger sequencing of the original plaque picked virus identified one extra deletion mutation in the L gene (15127-138), which would have led to a premature termination of the L protein. In NGS, the non-deleted sequence appeared to be present as a minor species (<5%). This appears to be caused by the accumulation of Defective Interfering (DI) particles which masked the standard virus sequence in Sanger sequencing. The absence of a type I interferon response in African Green Monkey kidney Vero cells allows the accumulation of defective interfering (DI) particles in MuV passages (Young *et al.*, 2009) which give rise to highly fluctuating titres in the passages, which were indeed observed (**Table S1**). In contrast, viruses populations passaged on B-LCL cells did not show their presence, as the extraordinary high coverage at the 3’ end of the antigenomic sequence (which represents the L gene and the region covered by DI particles) present in Vero cell passaged virus is absent in virus passaged on B-LCL cells (**Figure 1**). The accumulation of DI particles is also indicated by the substantial number of variant readings observed at the 3’end of the antigenome in virus populations passaged on Vero cells (see **Tables S2, S3**) and furthermore by the observation that in plaque assays the lowest dilution with the highest number of pfu did not show any plaques (**Table S1**) as at high concentrations DI particles prevent plaque formation.

Both PP1 and PP2 grew well and generated the same type of cytopathic effect in Vero cells (formation of syncytia) as the wild type MuV^G09^ non-recombinant progenitor virus isolated from the clinical material on Vero cells. In the case of PP2 these syncytia showed green fluorescence (**Fig 2**). The cytopathic effects of PP2 and MuV^009^ on B-LCL cells were the same and green fluorescence was observable in the clumps of the PP2 infected B-LCL cells floating in the medium.

### Assessment of the stability of the mutation in the clusters by NGS

Six additional passages of both viruses were carried at a high multiplicity of infection to provide maximum opportunity for the fixation of mutant genomes. PP1 was passaged on Vero cells and PP2 on B-LCL cells. As a control we also passaged the non-recombinant MuV^G09^ virus on Vero and B-LCL cells under the same conditions as PP1 and PP2. We chose these two cell substrates because of their different biological properties. Vero cells are adherent and give rise to syncytia. Vero cell passages of MuV were carried out at high MOI by infection of fresh Vero cell monolayers with supernatant virus from the previous passage. In the B-LCL cells, which are an IFN competent human B lymphocyte cell line that grow in suspension and leads to large cell clumps, we choose to allow maximum chances for the accumulation of mutations by carrying out the passages in such way that each passage represents an addition of fresh uninfected cells to the culture medium in which the cell clumps are dispersed by gentle shaking. The supernatant virus of each passage in B-LCL cells was titrated on Vero cells (**Table S1**).

The supernatant viruses from the Vero and B-LCL cell passages were pelleted by ultracentrifugation through a 25% (w/v) sucrose cushion and analysed by NGS on an Illumina platform with labelled primers so that the polarity of each read (positive or negative strand) could be determined.

Table 1 demonstrates that the constellation of the 11 mutations in the V/P gene of PP1 was maintained and stable over the 6 passages. The stability of PP1 in the passage series was not significantly different to that of the non-recombinant MuV^G09^ virus also passaged 6 times in parallel experiments on Vero cells. Similarly the 32 mutations in PP2 (Table 2) were also stable on passage in B-LCL cells and as stable as the wild type nucleotides at these positions in parallel passages of the MuV^G09^ virus. The frequencies with which the alternative readings occur in each cluster were in the order of 0.001 or 0.1% This in our experience is the normal frequency of alternative reads (0.04 to 0.10%) in NGS sequencing projects which may be generated during the amplifications involved in the library preparation and the sequence reading process itself. No significant predilection for changes that would restore the wild type MuV^G09^ nucleotide at any given position in the cluster of mutations was observed. The two other possible nucleotides at the mutated position were observed approximately twice as frequent (1.7-2.3) as those that would restore the original MuV^G09^ nucleotide at that position. This is what would be expected if the direction of variation was random rather than directed.

### Evolution of cluster-independent mutations during passage of rMuV

In order to ascertain that the passaging conditions did not impose some unexpected artefactual sequence stability we assessed whether mutations occurred during the passage series outside the clusters present in PP1 and PP2._ Many specific mutations accumulated to high frequencies during the six passages outside the clusters of originally mutated residues in PP1 and PP2. This indicates that though the virus could mutate in response to the changed cellular environment, the original sets of mutations were stably maintained. Examples of changes are given in Table 3. The more comprehensive representation of all the changes observed at a frequency of >1% in the deep sequencing reads are compiled and shown in **Table S2** and **S3** for PP1 and MuVG09 respectively.

**Table 3.**
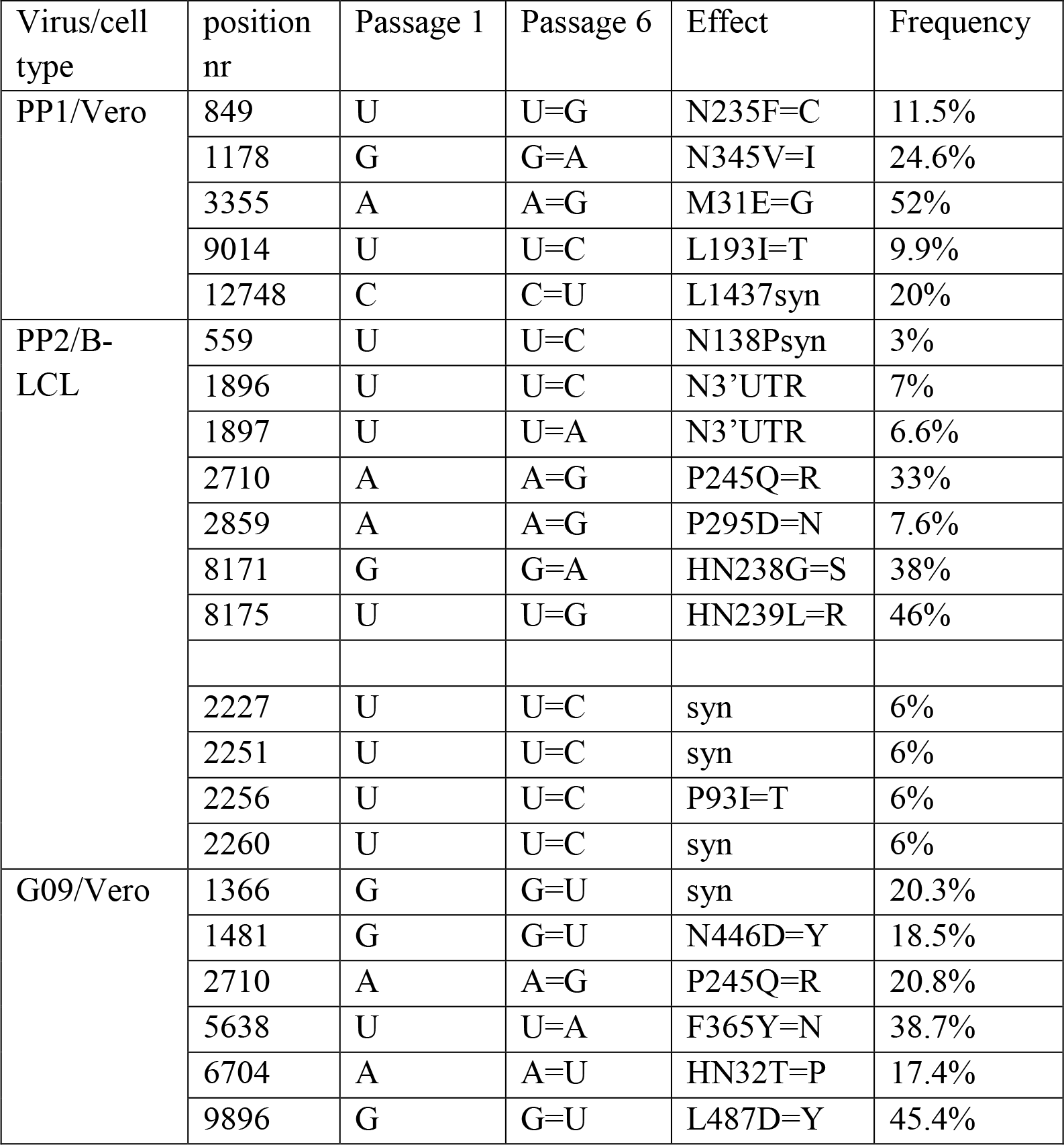
Examples of novel mutations observed after 6 passages of PP1, PP2 and the parent MuV^G09^ virus.

| Virus/cell type | position nr | Passage 1 | Passage 6 | Effect | Frequency |
| --- | --- | --- | --- | --- | --- |
| PP1/Vero | 849 | U | U=G | N235F=C | 11.5% |
|  | 1178 | G | G=A | N345V=I | 24.6% |
|  | 3355 | A | A=G | M31E=G | 52% |
|  | 9014 | U | U=C | L193I=T | 9.9% |
|  | 12748 | C | C=U | L1437syn | 20% |
| PP2/B-LCL | 559 | U | U=C | N138Psyn | 3% |
|  | 1896 | U | U=C | N3'UTR | 7% |
|  | 1897 | U | U=A | N3'UTR | 6.6% |
|  | 2710 | A | A=G | P245Q=R | 33% |
|  | 2859 | A | A=G | P295D=N | 7.6% |
|  | 8171 | G | G=A | HN238G=S | 38% |
|  | 8175 | U | U=G | HN239L=R | 46% |
|  | 2227 | U | U=C | syn | 6% |
|  | 2251 | U | U=C | syn | 6% |
|  | 2256 | U | U=C | P93I=T | 6% |
|  | 2260 | U | U=C | syn | 6% |
| G09/Vero | 1366 | G | G=U | syn | 20.3% |
|  | 1481 | G | G=U | N446D=Y | 18.5% |
|  | 2710 | A | A=G | P245Q=R | 20.8% |
|  | 5638 | U | U=A | F365Y=N | 38.7% |
|  | 6704 | A | A=U | HN32T=P | 17.4% |
|  | 9896 | G | G=U | L487D=Y | 45.4% |

Interestingly the NGS revealed that a number of mutations were present at a low frequency (~3.9%) in the fusion related external domain (FRED) of the F protein in both passage 1 and passage 6 of PP1 and of MuV^G09^. These were already present in passage 1, which is represents the 5^th^ passage in Vero cells after the original rescue and they were maintained at low frequency during the series. They affect the FRED domain by introduction of a number of charged residues that may well impact its functionality (Table 4). A similar observation was made in the NGS of the passages series of MuV^G09^ and thus this phenomenon is not specific to PP1 (data not shown).

**Table 4.** A cluster of mutations in the FRED of the fusion protein of PP1.

N terminus of F1 **FAGIAIGIAAALGVATAAQV**<sub>T</sub>\*
| Position | Mut | Effect | P1geno | P1 anti | P6c geno | P6canti |
| --- | --- | --- | --- | --- | --- | --- |
| 4869 | U=G | I=M | 3/385 | 4/58 | 1/176 | 3/28 |
| 4873 | A=U | I=F | 6/380 | 3/59 | 3/170 | 0/32 |
| 4881 | A=C | A syn | 3/365 | 4/59 | 3/170 | 3/35 |
| 4884 | C=G | L syn | 10/367 | 4/64 | 9/169 | 0/35 |
| 4888 | G=U | V=F | 5/360 | 2/66 | 4/169 | 1/36 |
| 4892 | C=A | A=E | 12/335 | 2/67 | 6/161 | 0/38 |
| 4894 | A=C | T=P | 19/328 | 9/65 | 13/159 | 4/38 |
| 4898 | C=A | A=E | 8/326 | 0/66 | 1/161 | 0/39 |
| 4901 | C=A | A=E | 12/330 | 1/66 | 5/161 | 1/39 |
| 4903 | C=A | Q=K | 22/329 | 7/66 | 11/159 | 1/38 |
| 4907 | U=A | V=E | 20/327 | 1/69 | 11/156 | 2/42 |
| total |  |  | 120/3832 | 37/705 | 67/1811 | 14/400 |
| frequency |  |  | 3.1% | 5.2% | 3.7% | 3.5% |

\* In large capitals the residues of the fusion related external domain that are affected by the mutations and in bold those that are changed as a result of a non-synonymous mutation.

In PP2 passage on B-LCL cells two variants seem to predominate with either a mutation at position 238 in the HN protein G > S or one at position 239 L > R. These are almost never present in the same RNA molecule and only a small number of original non-mutated wild type reads remain. The significance of the two mutations observed in the HN gene of PP2 during the passage series in B-LCL cells cannot easily be assessed because this is an unknown region of significance?

Noticeable is also that in PP2 between positions 2227 and 2260 in the sixth passage about 6 % of the reads showed a linked set of U to C mutations (Table 3) consistent with an interpretation that a number of biased hyper-mutated RNA molecules are carried along in passaged virus. One further interesting observation from these data is that unrelated to the passage series it is clear that PP1 but also other MuV viruses frequently insert extra nucleotides in the genomic sequences (**Table S4**). The significance of this is unclear. Similar observations have been made for other paramyxoviruses (our unpublished observations) such HPIV3, PIV5, hPIV2 and measles virus. In PP1 passaged on Vero cells these occur most frequently in the positions from about nt 12000 to the end of the (anti)genome representing most probable DI RNAs as they would likely affect the functionality of the L protein. These insertions occur primarily in two sets. The first is in the polyA sites. However the fact that the reads extend into the next gene indicates that these probably originate from read-through transcripts with extra As inserted. The G residue embedded in the homopolymeric stretch of A residues of the poly-adenylation signal (n.b. all expressed as + sequence) also is often observed to mutated to a G in these situations and these are also seen in the case of other virus in NGS studies. Both these types of changes may occur because during the generation of read-through transcripts the RdRp reads the stuttering signals normally associated with the generation of polyA tails of the mRNA. The second set of insertions at homopolymeric stretches in the area of the genome that is contained in DI RNA and in this case insertions of A, U and C nucleotides occur. The use of stuttering signals in the generation of Ebolavirus glycoprotein mutants during passage and reverse genetics has been well documented (Tsuda *et al.*, 2015). In the case described here however, the effect seems to be most frequently associated with read-through mRNA, the function of which remains unclear or potentially occurs in DI particles where function is no longer of importance.

### The EGFP gene does not vary to a greater extent than the virus genes

*A priori* the expectation was that the EFP gene in PP2 is not under selective constraint and that henceforth the ORF would have accumulated more mutations than the true virus genes. This appeared not to be the case when the number of variant readings at all positions in the ORFs encoding virus genes and that of EGFP were compared. The overall frequency of variant nucleotides at 0.20% was no greater in the EGFP ORF than in the other viral genes that presumably were under selective constraint (**Table S5**). This is an unexpected result but has been replicated in other paramyxoviruses with other fluorescent reporter proteins (our unpublished observations).

## Discussion

How and when during the rescue of PP1 and PP2 viruses the clusters of synonymous mutations arose is unclear. These replicating MuV were isolated as plaques in the wells in which the rescue experiment was performed. This is feasible because the rescue efficiency is relatively low and most wells in a six well plate do not contain more than one syncytium at 5-7 days post transfection. We have demonstrated here that once these clusters of mutations were generated the resulting replicator was stable. In the deep sequencing of viruses passaged 6 times at high multiplicity to allow for the maximum chance of the fixation of mutations no selection pressure to reversion was to the original nucleotide in the “wild type” sequence was observable. The direction of mutations appeared random and the variant readings were observed at such low frequencies as to probably be errors derived from the deep sequencing process rather than representing true variants in the virus population. It is also clear that the stability of the PP1 and PP2 replicating viruses did not reflect an inability of the viruses to fix mutations during these passage series as variations did occur during the passages in other genes and nucleotide positions in both the PP1 and PP2 viruses as well as in the parent virus upon passage. These were often found at very high frequencies (3-45%) even though many were non-synonymous. The stability of the clustered mutations thus indicates that they contribute to a stable and fit genotype that does not readily reverts back to the wild type constellations. Potential compensatory mutations in the V/P gene of PP1 and the N gene of PP2 were not observed as consistent features of the variations observed in NGS.

The stability of these mutant constellations is remarkable. As they consist primarily of synonymous mutations, it would be difficult to see a constraint at the protein coding level that would affect their reversion frequency. Non-synonymous mutations would probably be counter selected. The maintenance of the constellation in repeat passaging may point to a higher order RNA structural constraint but these would be predicted to be operative only at the mRNA level as the RNA in the + and − strand RNPs appears devoid of secondary structure in the paramyxoviruses (ref). Computational structure prediction of the mRNA did also not demonstrate a clear association with the generation of the clusters of mutations or in their maintenance (data not shown). The position of the mutant constellation is interesting. They are located at the 3’ end of the genome. This areas has been shown to be preferentially sensitive to biased hyper-mutation in measles virus (Otani *et al.*, 2014). Biased hyper-mutation is also prevalent in these MuV samples in the N gene and the start of the V/.P gene. The limited size of the constellations does also not affect the overall codon usage in these replicating MuV, which is known to be a specific feature of each paramyxovirus (Sciences and Ireland, no date) and the lack of synonymous mutations during viral evolution in the paramyxoviruses is observed but not explained.

How these mutant constellations were generated in the first place is an open question. The phenomenon described here may be specific for MuV. In our experience with rescue of other paramyxo- and pneumoviruses such as measles, canine distemper and rinderpest viruses - as well as respiratory syncytial virus (Gassen *et al.*, 2000; Moeller *et al.*, 2001; Brown *et al.*, 2005; Lemon *et al.*, 2015) - we have not encountered this phenomenon apart from occasional clusters of mutation that were generated by biased hypermutation involving primarily U to C and at a lesser frequency A to G changes. The clusters of mutations in the PP1 and PP2 viruses do not show this bias. The value for κ i.e. the ratio of transitions over transversions in all clusters summed together was 4.9 (bias towards transitions) which is similar to that found in between genotype comparisons for MuV. It is not due to the presence of the extra EGFP gene as it occurred both in PP1 and PP2.

It seems unlikely that the limited number of replications required to generate a syncytium during the primary isolation of these viruses would allow sequential selection of the large number of mutations that would provide this stable constellation. Furthermore, the observation that these mutations are clustered indicates a different mechanism for their generation. We suggest that the most likely process that generated the clusters is the transcription of the DNA plasmid by T7 RNA polymerase followed by removal of unfit viruses by selection against viruses with lethal mutations in the N and V/P ORFs. Why the mutations occur in clusters remain an open question. One possible explanation for the occurrence of the clusters of mutation may be that quantum biological effects led to entanglement of the protons in what is essentially a proton code in the plasmid DNA formed by the patterns of two (A-U) or three (G-C) hydrogen bonds read by the T7 RNA polymerase. Entanglement has been invoked in a number of studies dealing with mutations in biological systems (Pusuluk and Deliduman, 2010)(Al-khalili, 2013) as wells as in catalysis of ATP independent cleavage of DNA by restriction endonucleases (Kurian, Dunston and Lindesay, 2014). On the basis of modelling studies Rieper et al. (Rieper, Anders and Vedral, 2010) suggested that nucleotides in DNA might be read in the context of their neighbouring nucleotides, which may explain constraints on synonymous mutation in RNA viruses (Rima, 2015). However, whilst quantum biological effects merit more attention in virology, their experimental verification remains a challenge (Al-khalili, 2013).

### Author statements

The study was funded by Queen’s University Belfast and St Andrews University. The authors declare no conflicts of interest. Ethical requirements have been fulfilled as no animal or human subjects were used.

## Materials and Methods

### Viruses and Cells

Clinical material was obtained from Dr Paul Rota (CDC Atlanta) as a buccal swab from a patient during the 2009/10 US outbreak of mumps in New York caused by a genotype G5 virus (Xu *et al.*, 2011). We isolated the virus on Vero cells, passaged four times in Vero cells and named it here abbreviated to MuV^G09^. The complete consensus sequences of the viral RNA in the tissue sample and the isolated viruses were determined by classical Sanger sequencing of overlapping RT-PCR amplicons (sequences of primers available on request) and found to be identical.

Vero cells were used for isolation, rescue and routine passage of the viruses as well as titration of plaque forming units (pfu). B-LCL cells were obtained from Erasmus University Medical Centre, Rotterdam

### Generation of recombinant viruses

Plasmids that expressed the N, V/P and L proteins with the authentic sequence of the MuV^G09^ virus were generated to act as helper plasmids in the rescue experiments. Plasmids were also generated that represented the full length consensus sequence of the viral RNA which was identical to that of the isolated virus MuV^G09^ as well as one in which the enhanced green fluorescent protein gene (EGFP) was inserted as an additional transcription unit between the V/P gene and the M gene of MuV. These sequences were placed between a T7 promotor and hepatitis delta ribozyme. After infection of Vero cells with Fowl Pox-expressing T7, the cell monolayer was transfected with the helper and full lengths cDNA plasmids and the appearance of syncytia was monitored over 1 week. Usually never more than one syncytium was observed per well. These were aspirated and propagated further on Vero cells for 4 low MOI passages.

### Next generation Sequencing (NGS)

The RNA was extracted from infected cells and subjected to total RNA with ribosomal and mitochondrial RNA reduction library preparation as per manufacturer’s instructions (Illumina). The samples were sequenced using the Hi-seq illuminated platform. The sequencing data was subjected to directional analysis which separates the reads based on directionality allowing the isolation of the viral genome (negative sense) and viral mRNA reads (positive sense). The isolated viral genome reads were then aligned to the MuV reference sequence using BWA alignment software. An in-house script was then used to enumerate the SNPs at each nucleotide of the reference sequence.

Figure 1 legend: **Coverage of reads and preponderance of Defective Interfering RNAs in passages of viruses on Vero and B-LCL cells** Diagrams that show the total numbers of reads obtained in NGS for each nucleotide over the entire mumps genome of 15372 nucleotides displayed by the Tablet programme (Milne *et al.*, 2013). In Fig 1A the scale is 1-3000 in Fig 1B it is 1-1,500,000 indicating the extraordinary accumulation of reads associated with the appearance of DI particles.

Figure 2 legend: **Rescue of the viruses as replicators; cpe and fluorescent plaques.**

Figure 2A: Generation of rMuV^G09^ by reverse genetics.

Panel 1 shows the presence of primary foci of rescue at 5 days post transfection; panel 2 shows primary syncytia which were plaque picked and subsequently Vero cells were infected with the aspirated virus stocks. CPE was detected 1-2 dpi. Panel 3 shows plaque picked rMuV^G09^ grown for 4 low MOI passages on Vero cells. All show characteristic syncytium-formation.

Figure 2 B: Generation of rMuV^G09^ expressing EGFP - rMuV^G09^EGFP(3) - by reverse genetics.

Panel 1 shows the presence of primary foci of rescue at 5 days post transfection in both phase contrast and UV microscopy; panel 2 shows primary syncytia which were plaque picked and subsequently Vero cells were infected with the aspirated virus stocks. EGFP expression was evident 1 dpi and cpe was detected 1-2 dpi. Panel 3 shows plaque picked rMuV^G09^EGFP(3) grown for 4 low MOI passages on Vero cells Passaged virus images show characteristic syncytium-formation.

